# A frequency-domain machine learning method for dual-calibrated fMRI mapping of oxygen extraction fraction (OEF) and cerebral metabolic rate of oxygen consumption (CMRO_2_)

**DOI:** 10.1101/660357

**Authors:** Michael Germuska, Hannah Chandler, Thomas Okell, Fabrizio Fasano, Valentina Tomassini, Kevin Murphy, Richard Wise

**Affiliations:** CUBRIC, Department of Psychology, Cardiff University, Cardiff, United Kingdom; Wellcome Centre for Integrative Neuroimaging, FMRIB, Nuffield Department of Clinical Neurosciences, University of Oxford, United Kingdom; Siemens Healthcare Ltd, Frimley, Camberley, UK; Division of Psychological Medicine and Clinical Neurosciences, Cardiff University School of Medicine, Cardiff, United Kingdom; Department of Neuroscience, Imaging and Clinical Sciences, “G. D’Annunzio University” of Chieti-Pescara, 66100, Chieti, Italy; Institute for Advanced Biomedical Technologies, “G. D’Annunzio University” of Chieti-Pescara, 66100, Chieti, Italy

**Keywords:** magnetic resonance imaging, metabolism, oxygen extraction fraction, CMRO_2_, OEF, BOLD, artificial neural networks, machine learning, calibrated-fMRI

## Abstract

Magnetic resonance imaging (MRI) offers the possibility to non-invasively map the brain’s metabolic oxygen consumption (CMRO_2_), which is essential for understanding and monitoring neural function in both health and disease. However, in depth study of oxygen metabolism with MRI has so far been hindered by the lack of robust methods. One MRI method of mapping CMRO_2_ is based on the simultaneous acquisition of cerebral blood flow (CBF) and blood oxygen level dependent (BOLD) weighted images during respiratory modulation of both oxygen and carbon dioxide. Although this dual-calibrated methodology has shown promise in the research setting, current analysis methods are unstable in the presence of noise and/or are computationally demanding. In this paper, we present a machine learning implementation for the multi-parametric assessment of dual-calibrated fMRI data. The proposed method aims to address the issues of stability, accuracy, and computational overhead, removing significant barriers to the investigation of oxygen metabolism with MRI. The method utilizes a time-frequency transformation of the acquired perfusion and BOLD-weighted data, from which appropriate feature vectors are selected for training of machine learning regressors. The implemented machine learning methods are chosen for their robustness to noise and their ability to map complex non-linear relationships (such as those that exist between BOLD signal weighting and blood oxygenation). An extremely randomized trees (ET) regressor is used to estimate resting blood flow and a multi-layer perceptron (MLP) is used to estimate CMRO_2_ and the oxygen extraction fraction (OEF). Synthetic data with additive noise are used to train the regressors, with data simulated to cover a wide range of physiologically plausible parameters. The performance of the implemented analysis method is compared to published methods both in simulation and with *in-vivo* data (n=30). The proposed method is demonstrated to significantly reduce computation time, error, and proportional bias in both CMRO_2_ and OEF estimates. The introduction of the proposed analysis pipeline has the potential to not only increase the detectability of metabolic difference between groups of subjects, but may also allow for single subject examinations within a clinical context.

## 1 Introduction

Under normal conditions the brain’s energy needs are met via a continuous supply of oxygen and glucose for the local production of ATP via aerobic metabolism (Verweij et al., 2007). Any disruption of the supply of oxygen to the brain tissue can have significant consequences (Safar, 1988), and impaired cerebral oxygen metabolism is associated with a wide variety of neurological conditions (Frackowiak et al., 1988; Ishii et al., 1996; Miles and Williams, 2008). Therefore, monitoring and mapping the brain’s consumption of oxygen is vital for understanding the diseases and mechanisms by which the metabolic consumption of oxygen may be affected. The cerebral metabolic rate of oxygen consumption (CMRO_2_) has traditionally been measured with positron emission tomography (Frackowiak et al., 1980). However, this method has some substantial limitations including the use of ionizing radiation and the need for local production of 15-oxygen labeled tracers. Due to these limitations there is great interest in developing alternative, non-invasive, methods of mapping CMRO_2_. One promising technique of non-invasively mapping CMRO_2_ is the so-called dual-calibrated fMRI (dc-fMRI) method (Bulte et al., 2012; Gauthier et al., 2012). This method is finding growing adoption in the research setting, and has already been applied in Alzheimer’s disease (Lajoie et al., 2017), carotid artery occlusion (De Vis et al., 2015), and studies of pharmacological modulation (Merola et al., 2017). For a review of the method and details on the its practical application please see (Germuska and Wise, 2019). Despite the promise shown by this technique, the reported between-session repeatability is relatively low (Merola et al., 2018) and improvements in the data acquisition and/or analysis are required if individualized assessment is to be made possible.

One of the key difficulties in analyzing dual-calibrated fMRI data is noise propagation through the analysis pipeline, which leads to unstable parameter estimates. We have previously presented regularized non-linear least squares fitting approaches that utilize prior physiological knowledge to produce more robust parameter estimates (Germuska et al., 2019; Germuska et al., 2016). Even though such regularization reduces the mean square error it does so by trading off a reduction in variance with an increase in bias. An alternative approach to reduce the prediction error is the use of noise insensitive machine learning regression methods. Decision tree based regression methods, for example random forest (Breiman, 2001) and extremely randomized trees (Geurts et al., 2006), are robust to both output (Breiman, 2001; Geurts et al., 2006) and input noise (Yue et al., 2018) and are able to capture non-linear relationships between input features and target parameters. This noise immunity is likely due to the randomization included in the choices of features at splitting nodes (random forest) and cut-points (extremely randomized trees), which improve the generalizability of the regressors. For non-linear mappings with a high degree of complexity artificial neural networks such as the multi-layer perceptron (MLP), a feedforward network with multiple hidden layers, offer a machine learning method that is inherently robust to noise (Bernier et al., 1999). In this paper we present an analysis pipeline comprised of an extremely randomized trees regressor and a MLP, cascaded to infer resting CBF and CMRO_2_ from dual-calibrated fMRI data. A frequency-domain representation of simulated MRI data with the additive noise is used to train each of the regressors. Simulated data has the advantage over *in-vivo* data in this application as it allows a balanced dataset to be generated that covers a broad range of physiological variation. Such a dataset is essential to avoid bias in parameter estimation and to provide generalizability across groups and diseases. A frequency-domain representation is chosen as it allows for convenient dimensionality reduction, with most of the information of interest encoded at low temporal frequencies, and takes advantage of the superior ability of artificial neural networks to learn discriminative features from frequency-domain representation of a signal compared to a time-domain representation (Hertel et al., 2016). The performance of the proposed machine learning (ML) implementation is compared to an existing regularized non-linear least squares (rNLS) method (Germuska et al., 2019) both in simulation and in data acquired from a cohort of 30 healthy volunteers. We hypothesized that the machine learning approach would be able to achieve comparable or reduced prediction error with significantly reduced bias and computational overhead.

## 2 MRI Data Acquisition

Thirty healthy volunteers (16 males, mean age 32.53 ± 6.06 years) were recruited to the study. The local ethics committee approved the study and written informed consent was obtained from each participant. Blood samples were drawn via a finger prick prior to scanning and were analyzed with the HemoCue Hb 301 System (HemoCue, Ängelholm, Sweden) to calculate the systemic [Hb] value for each participant. All data was acquired using a Siemens MAGNETOM Prisma (Siemens Healthcare GmbH, Erlangen) 3T clinical scanner with a 32-channel receiver head coil (Siemens Healthcare GmbH, Erlangen). The acquisition protocol was as previously described (Germuska et al., 2019). Briefly, an 18-minute dual-excitation pseudo-continuous arterial spin labeling (pCASL) and BOLD-weighted acquisition was acquired during modulation of inspired oxygen and carbon dioxide. Gas modulation was performed according to a protocol previously proposed by our lab (Germuska et al., 2016), and end-tidal monitoring was performed throughout the acquisition from the volunteer’s facemask using a rapidly responding gas analyzer (PowerLab®, ADInstruments, Sydney, Australia). The prototype pCASL sequence (Germuska et al., 2019) parameters were as follows: post-labeling delay and label duration 1.5 seconds, EPI readout with GRAPPA acceleration (factor = 3), TE_1_ = 10ms, TE_2_ = 30ms, TR = 4.4 seconds, 3.4 × 3.4mm in-plane resolution, and 15 (7mm) slices with 20% slice gap.

## 3 Synthetic MRI Data Generation

Synthetic data was simulated to match the 18-minute *in-vivo* acquisition protocol using standard physiological models for the change in BOLD signal (Bulte et al., 2012; Gauthier and Hoge, 2013; Wise et al., 2013), as summarized by equation 1.

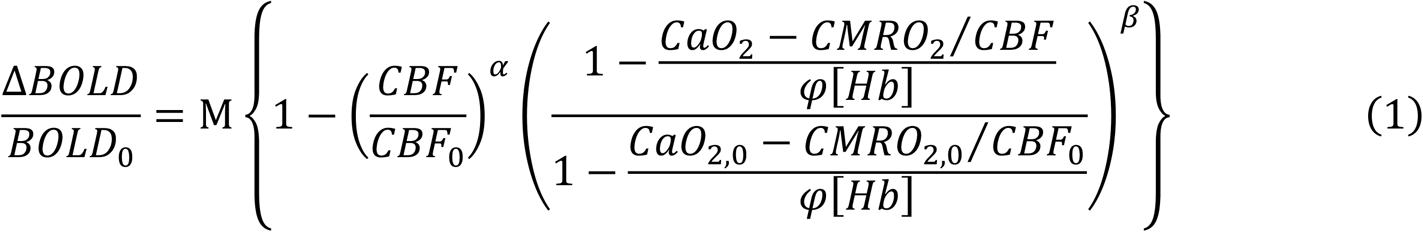

Where, Δ*BOLD*/*BOLD*_0_ is the fractional change in BOLD signal due to a change in arterial oxygen content (CaO_2_) or CBF due to either a hyperoxic or hypercapnic respiratory stimulus. M is a lumped parameter that is equal to *K* · ((1 − *SvO*_2_)·[*Hb*])^*β*^. Where K is a scaling factor dependent on the field strength, resting venous blood volume, tissue structure, and water diffusion effects in the extravascular space. [Hb] is the blood hemoglobin concentration and SvO_2_ is the venous oxygen saturation. φ is the oxygen binding capacity for Hb (1.34 ml/g), α is the Grubb exponent that couples blood volume and blood flow changes, and β is a field strength dependent constant that summarizes the non-linear effects associated with the tissue structure and water diffusion effects. The values of α and β were fixed to the optimized values (0.06 and 1) found by (Merola et al., 2016), which minimize the error in OEF estimates over a range of vascular physiology. The subscript 0 represents the baseline or resting state. The hyperoxic and hypercapnic stimuli are assumed to be iso-metabolic, so CMRO_2_ = CMRO_2,0_.

The arterial spin labeling signal was modeled according to the simplified pCASL kinetic model (Alsop et al., 2015), and physiological constraints on baseline parameters were applied according to a simple model of oxygen exchange (Gjedde, 2002; Hayashi et al., 2003), equation 2.

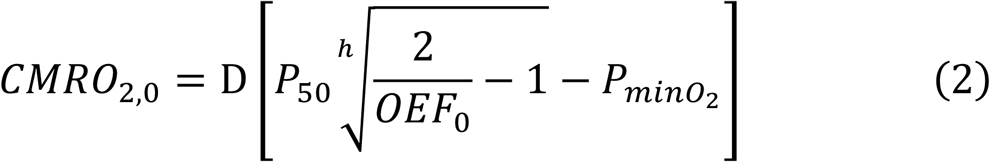

Where D is the effective oxygen diffusivity of the capillary network and can be expressed as a product of the effective oxygen permeability and the capillary blood volume, *D* = *κ* · *CBV*_*cap*_. P_50_ is the blood oxygen tension at which hemoglobin is 50% saturated (26 mmHg), h is the Hill coefficient (2.8) and 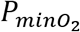 is the minimum oxygen tension at the mitochondria (which is thought to be negligible in healthy tissue (Gjedde, 2002)). In the modeling we assume a fixed value for κ of 3 μmol/mmHg/ml/min, corresponding to a typical diffusivity of 3 (Mintun et al., 2001) to 4 μmol/100g/mmHg/min (Vafaee and Gjedde, 2000) for CBV_cap_ = 1 to 1.33 ml/100g. The physiological parameter space encompasses a wide range of plausible physiology including both healthy and dysfunctional brain tissue, and is summarized in Table 1. A summary of MRI abbreviations and all model parameters used in the simulations is given in table 2.

**Table 1.**
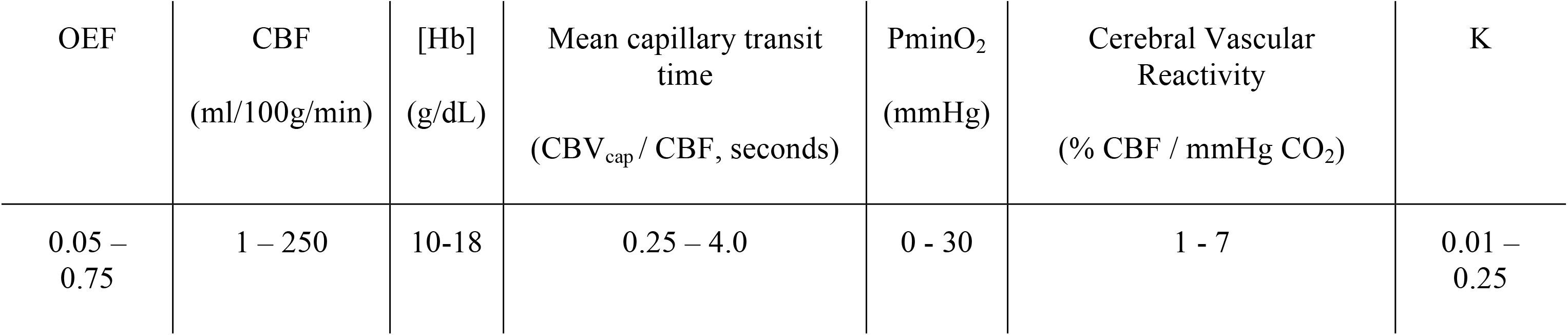
Range of physiological parameters used in the dc-fMRI data simulations for training of the machine learning regressors.

| OEF | CBF<br>(ml/100g/min) | [Hb]<br>(g/dL) | Mean capillary transit<br>time<br>(CBV<sub>cap</sub> / CBF, seconds) | PminO<sub>2</sub><br>(mmHg) | Cerebral Vascular<br>Reactivity<br>(% CBF / mmHg CO<sub>2</sub>) | K |
| --- | --- | --- | --- | --- | --- | --- |
| 0.05 –<br>0.75 | 1 – 250 | 10-18 | 0.25 – 4.0 | 0 - 30 | 1 - 7 | 0.01 –<br>0.25 |

**Table 2.**
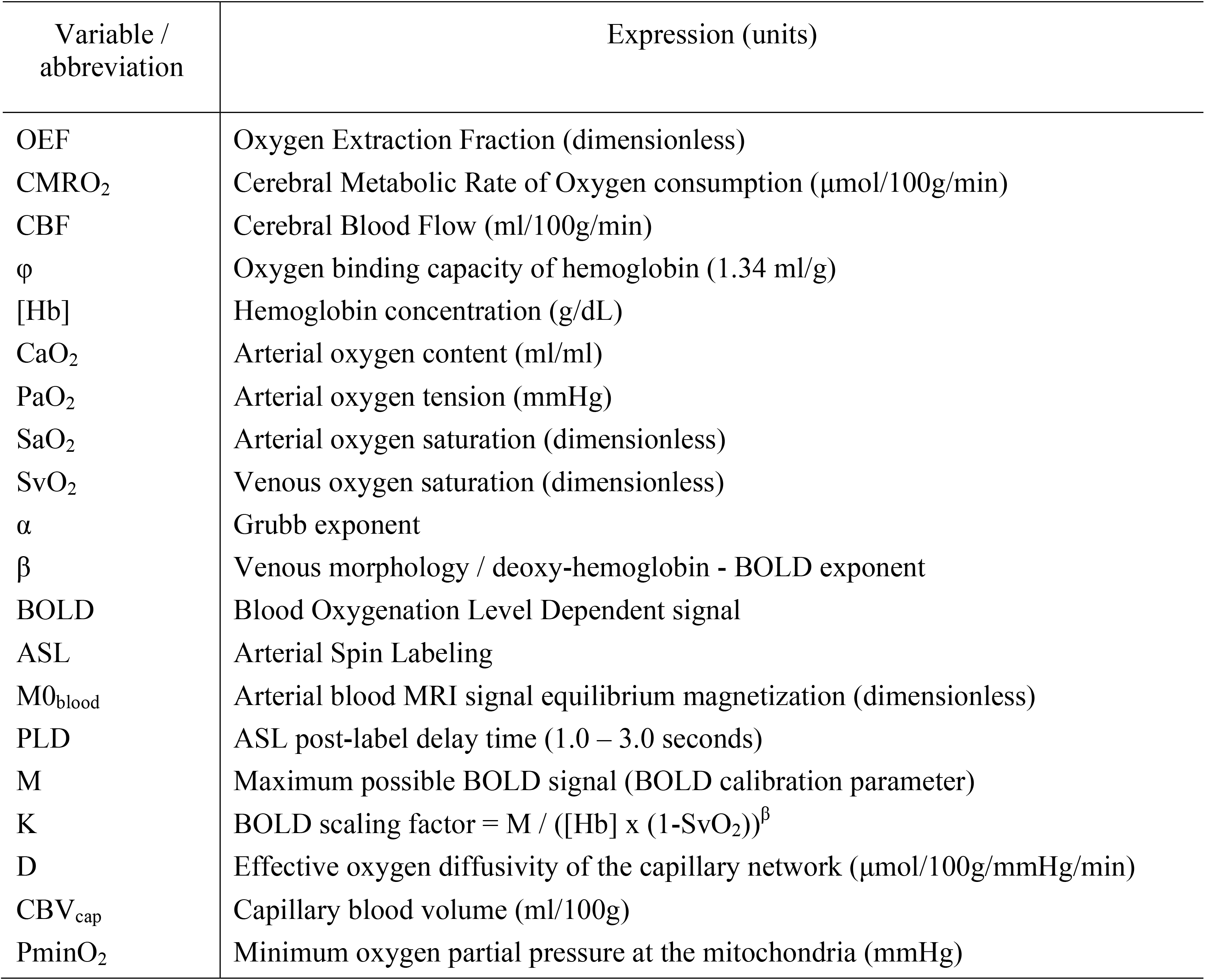
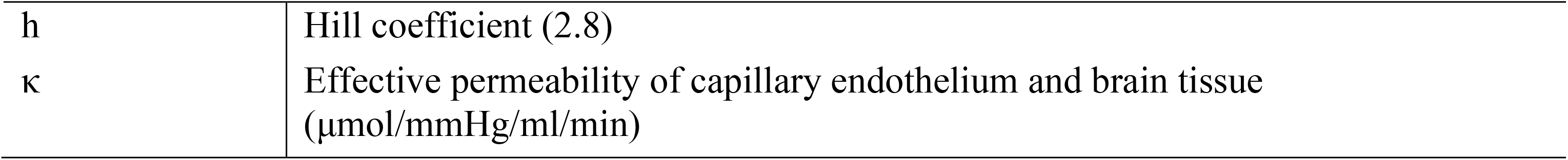
Summary of model parameters and abbreviation used in the dc-fMRI data simulations and their definitions.

| Variable /<br>abbreviation | Expression (units) |
| --- | --- |
| OEF | Oxygen Extraction Fraction (dimensionless) |
| CMRO<sub>2</sub> | Cerebral Metabolic Rate of Oxygen consumption (μmol/100g/min) |
| CBF | Cerebral Blood Flow (ml/100g/min) |
| φ | Oxygen binding capacity of hemoglobin (1.34 ml/g) |
| [Hb] | Hemoglobin concentration (g/dL) |
| CaO<sub>2</sub> | Arterial oxygen content (ml/ml) |
| PaO<sub>2</sub> | Arterial oxygen tension (mmHg) |
| SaO<sub>2</sub> | Arterial oxygen saturation (dimensionless) |
| SvO<sub>2</sub> | Venous oxygen saturation (dimensionless) |
| α | Grubb exponent |
| β | Venous morphology / deoxy-hemoglobin - BOLD exponent |
| BOLD | Blood Oxygenation Level Dependent signal |
| ASL | Arterial Spin Labeling |
| M<sub>0blood</sub> | Arterial blood MRI signal equilibrium magnetization (dimensionless) |
| PLD | ASL post-label delay time (1.0 – 3.0 seconds) |
| M | Maximum possible BOLD signal (BOLD calibration parameter) |
| K | BOLD scaling factor = $M / ([Hb] \times (1 - SvO_2))^\beta$ |
| D | Effective oxygen diffusivity of the capillary network (μmol/100g/mmHg/min) |
| CBV<sub>cap</sub> | Capillary blood volume (ml/100g) |
| PminO<sub>2</sub> | Minimum oxygen partial pressure at the mitochondria (mmHg) |
| h | Hill coefficient (2.8) |
| $\kappa$ | Effective permeability of capillary endothelium and brain tissue ($\mu\text{mol}/\text{mmHg}/\text{ml}/\text{min}$) |

The partial pressure of arterial oxygen (PaO_2_) and change in carbon dioxide (ΔPaCO_2_) were modeled to match the range of end-tidal recordings acquired from healthy volunteers. The baseline PaO_2_ had a range of 90-120 mmHg, ΔPaO_2_ was 200 to 300 mmHg, and ΔPaCO_2_ was set to 8-12 mmHg. Rectangular stimulus blocks were convolved with a gamma density function with shape parameter 0.5-2.5 to account for the variation in biological rise and fall times of the hyperoxic and hypercapnic stimuli. Drift in ΔPaCO_2_, which was observed in some subjects, was included by adding a bandpass filtered noise signal (4^th^ order IIR filter, lowcut/highcut = 0.005/0.05 of the Nyquist frequency). Change in the arterial blood longitudinal relaxation rate due to dissolved oxygen was included in pCASL calculations as per (Germuska et al., 2019). Noise (BOLD tSNR = 90, pCASL tSNR = 3 for CBF = 60 ml/100g/min) was added to simulated BOLD and pCASL time series. The pCASL noise was bandpass filtered (4^th^ order IIR filter, lowcut/highcut = 0.05/0.8 of the Nyquist frequency) and the BOLD noise was lowpass filtered (1^st^ order IIR filter, highcut = 0.5 of the Nyquist frequency) to match the noise characteristics of the *in-vivo* data. In addition, the BOLD timeseries data was highpass filtered with a 320 second cut-off using the filter implementation in FSL (Jenkinson et al., 2012), which is routinely used for de-trending fMRI data. Figure 1 shows 50 randomly generated pCASL and BOLD timeseries overlaid with the temporal mean to demonstrate the typical output of the simulations. Please note that the pCASL timeseries are divided by the equilibrium magnetization of arterial blood (M0_blood_), and the baseline signal has been set to zero for display purposes.

**Figure 1.**
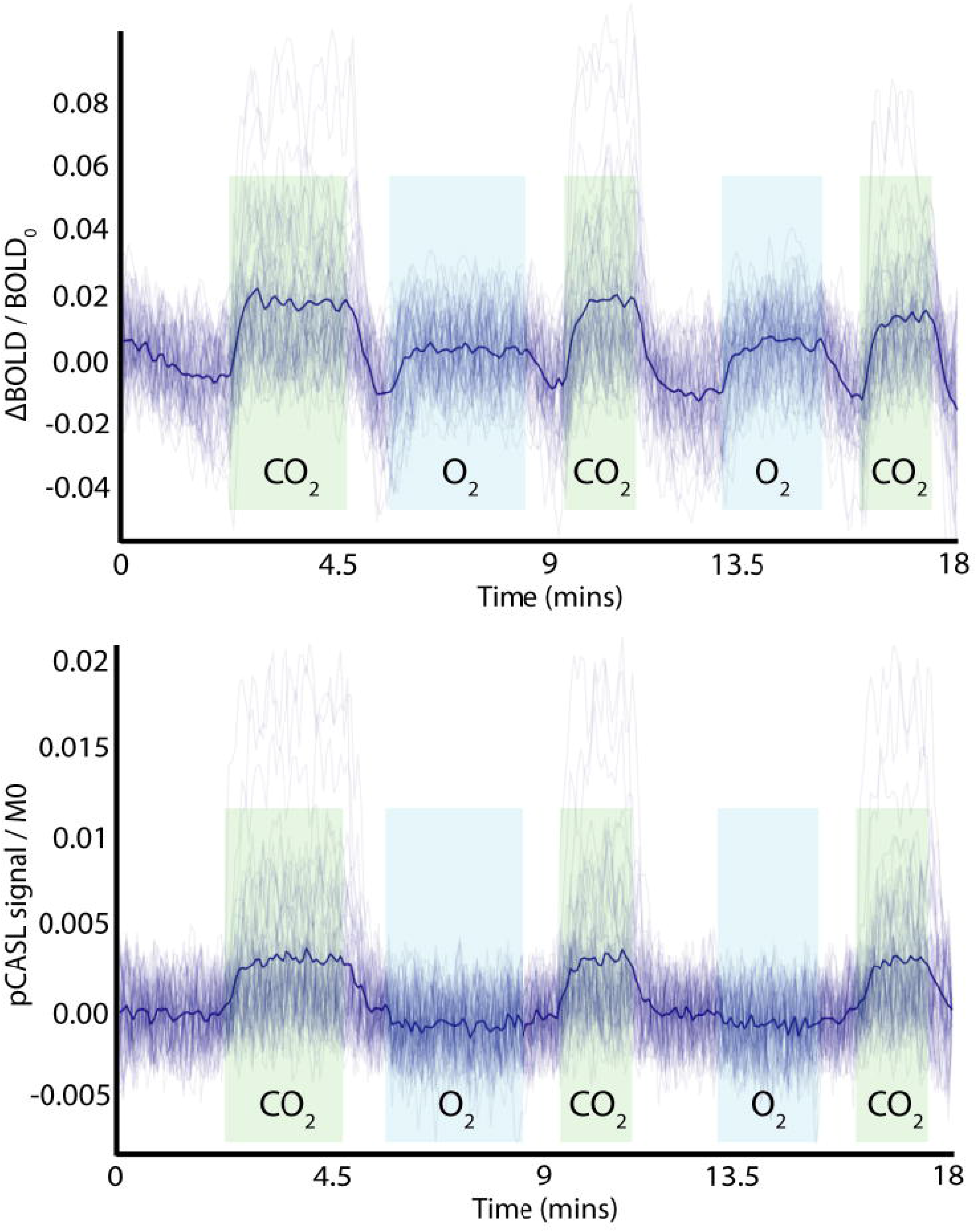
Example of simulated time-domain data (BOLD and ASL) with added noise and variation in physiological parameters, showing periods of hypercapnic (green) and hyperoxic (light blue) stimuli. The dark blue line represents the mean time-course over the example time series. Note the pCASL signal is normalised by the equilibrium magnetisation of arterial blood (MO) and has the baseline signal subtracted for display purposes.

## 4 Methods

A schematic diagram describing the analysis/training pipeline is shown in figure 2. ASL and BOLD timeseries data, either simulated (as described in section 3) or *in-vivo* data, are Fourier transformed into magnitude and phase data. This frequency domain data is then truncated after the first 15 data points (low pass filtered) and combined with physiological recordings and sequence parameters to create a feature vector for model training/prediction (if *in-vivo* data is being analyzed). Parameter estimation is carried out in a two-stage process; first the resting blood flow (CBF_0_) is estimated, and then rate of oxygen consumption.

**Figure 2.**
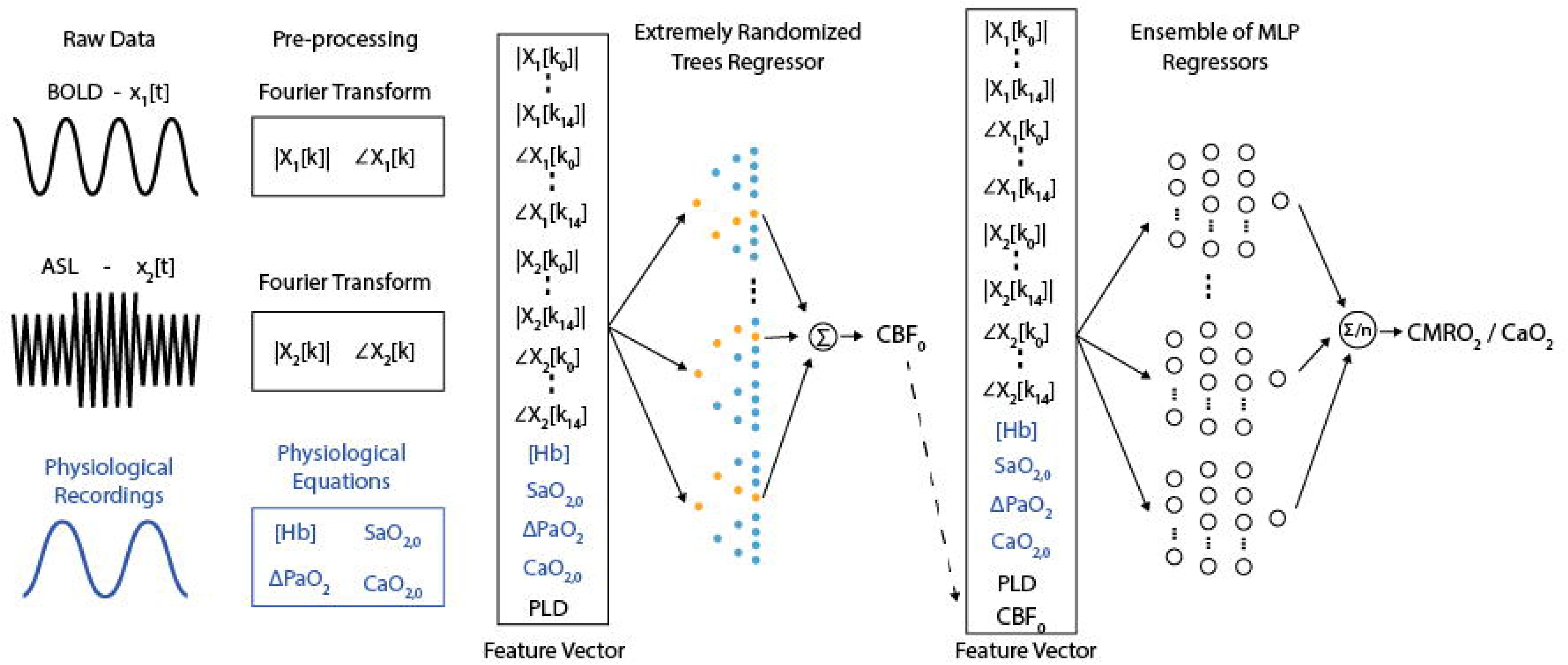
Schematic diagram of the frequency-domain machine learning pipeline. Raw data is pre-processed prior to the construction of a feature vector. This initial feature vector is used to estimate baseline perfusion. The perfusion estimate is then included in the feature vector fed into an ensemble of multilayer perceptron networks used to estimate the resting rate of oxygen metabolism.

Truncation of the frequency domain data removes high-frequency content that is unrelated to either the hyperoxic or hypercapnic respiratory modulations and thus removes noise from the training data. The resting blood flow is estimated separately from the rate of oxygen consumption to reduce the complexity of the required mapping between the MRI data and the target parameters. Additionally, the use of extremely randomized trees (ET) regression rather than an artificial neural network at this stage in the pipeline takes full advantage of the noise immunity of decision tree based methods (Yue et al., 2018) and reduces the potential of overfitting. The inclusion of the post-label delay in the feature vector is necessary to incorporate an implicit slice timing correction for CBF_0_ calculation, while the blood oxygenation parameters ([Hb], ΔPaO_2_, SaO_2,0_, CaO_2,0_) are included here due to the influence of dissolved oxygen on the longitudinal relaxation rate of arterial blood. In total each feature vector that is input into the ET regressor consists of 65 entries.

The result of the ET regression is then incorporated into the feature vector (now 66 entries) and input into an ensemble of MLPs to predict CMRO_2,0_ / CaO_2,0_, from which CMRO_2,0_ and OEF_0_ can be calculated (CMRO_2_ / CaO_2_ = OEF × CBF via the Fick principle). The blood oxygenation parameters in this case not only inform on the relaxation rate of arterial blood, but also link the CBF and BOLD signal changes to the underlying metabolic parameters as described by equation 1. In practice each MLP in the ensemble is trained individually, with the average of their predictions being used for inference when deployed for the analysis of *in-vivo* data.

The ET regressor and MLP were implemented in Scikit learn (Pedregosa et al., 2011). The extremely randomized trees regressor was trained with the following options, number of estimators = 50, bootstrap = True, and out-of-bag samples were used to estimate the R^2^ on unseen data. A total of 50,000 simulations were used for training. The MLP network has two-hidden layers and 50 nodes in each layer. The activation function for each node was chosen to be a rectified linear unit (ReLU). The ADAM solver was used for training with 1×10^6^ simulated feature vectors and 10% of the data were used for early stopping. Data simulation and training was repeated 40 times to create an ensemble of MLP networks to further reduce the uncertainty in parameter estimates (Sollich and Krogh, 1996).

The validation score for the extremely randomized trees regressor for predicting resting cerebral blood flow was 0.997, slightly greater than the results obtained for a random forest implementation (0.961). The validation score for the MLP estimation of CMRO_2,0_ / CaO_2,0_ were 0.923 ± 0.002. Training of the MLP network was also undertaken while eliminating key elements of the simulation or feature vectors to see how this affected the performance of the MLP. When BOLD data was excluded from the feature vector the validation score dropped to 0.577. Excluding the CO_2_ and O_2_ stimuli (but including the BOLD data) reduced the validation scores to 0.63 and 0.71 respectively.

A further 5,000 simulated datasets (with OEF restricted to 0.15 to 0.65, all other parameters as in table 1) were constructed to compare the performance of the proposed machine learning implementation with a previously implemented regularized non-linear least squares fitting method (Germuska et al., 2019). Each method was compared to the simulated data using a robust regression method (bisquare) in terms of the RMS error and proportional bias. A bisquare cost function was used for the regression to reduce the influence of outliers and allow a robust estimate of the proportional bias. The rNLS fitting was implemented with regularization applied to the resting OEF and the effective oxygen diffusivity (D), as previously described. The relative weighting between OEF and diffusivity regularization was maintained constant, as per the optimization in (Germuska et al., 2019). However, the total weighting was varied to assess the impact on OEF and CMRO_2_ error and proportional bias (slope of the simulated parameter values plotted against the parameter estimates).

## 5 Results

### 5.1 Simulations

Analysis of the simulated data demonstrated a substantial reduction in the RMS error of machine learning OEF estimates compared to rNLS estimates. The bisquare RMS error was 0.047 when using the mean prediction from the 40 MLP networks, and 0.055 for a randomly chosen MLP network. The rNLS approach produced a minimum bisquare RMS error of 0.094. The ML approach displayed negligible proportional bias in OEF estimates (slope of true vs. estimated values = 0.982), whereas rNLS estimates had variable levels of bias depending on the level of regularization, see figure 3a for a summary of the results. As expected from the OEF results, ML estimates of CMRO_2_ also had significantly reduced error and bias compared to the rNLS implementation. The proportional bias for the ML implementation was 0.977 compared to a minimum bias of 0.913 for the rNLS method. The bisquare RMS error in CMRO_2_ estimates for the ML implementation was 20.3 μmol/100g/min (22.6 for an individual MLP network) whereas the error for rNLS estimates ranged from 29.6 to 52.4 μmol/100g/min depending on the level of bias (with greater bias coinciding with reduced error), see figure 3b.

**Figure 3.**
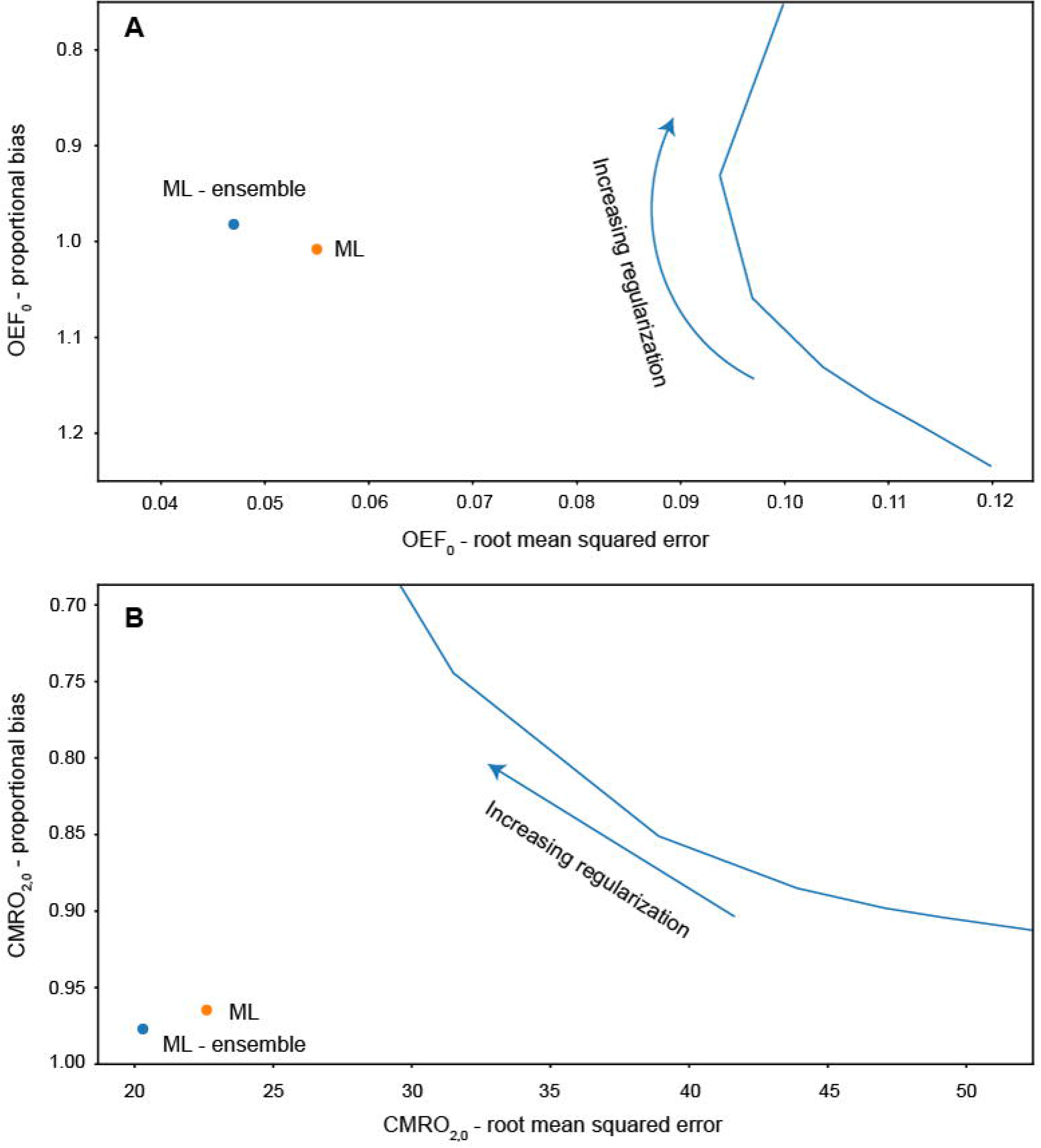
Root mean squared error and proportional bias in OEF_0_ (A) and CMRO_2,0_ (B) estimates for each analysis method fitting to simulated data (5000 simulations). Solid blue line plots the error and bias for increasing regularization weighting for the regularized non-linear least squares analysis

Training of the MLP with reduced feature vectors (excluding the BOLD data) or limited respiratory stimuli (excluding either CO_2_ or O_2_ modulation) highlights the importance of each signal and stimulus in estimate the rate of oxygen consumption. As expected, removing the BOLD signal resulted in a significant reduction in the network’s ability to estimate CMRO_2_ (validation R^2^ reduced from 0.923 for the full model to 0.58). In this instance there should be no information relating to OEF in the feature vector and so the inference is based solely on the correlation between baseline flow and CMRO_2_ in the simulated data. Adding the BOLD data back in but with only an O_2_ stimulus does little to improve the performance of the network (R^2^ = 0.63). This is not unexpected as the hyperoxic BOLD signal is largely related to venous blood volume (Blockley et al., 2013) with little influence from OEF. Perhaps unexpectedly, including the CO_2_ stimulus but not the O_2_ stimulus significantly improves the ability of the network to infer resting CMRO_2_ (R^2^ = 0.71). While this is still significantly worse than the full model, it suggests that some quantitative metabolic information may be extracted from hypercapnic calibration studies that are normally employed to estimate relative changes in CMRO_2_ (Hoge, 2012). Additionally, such results suggest that the simulation framework could be utilized to optimize data acquisition by designing respiratory stimuli that maximize the performance of the ML implementation, and that such respiratory paradigms may be different compared to those for standard analysis methods (which are unable to infer resting CMRO_2_ information from a hypercapnic calibration experiment).

### 5.2 In-vivo

Due to the limited availability and technical challenges associated with acquiring 15-oxygen PET data for CMRO_2_ mapping (the gold standard approach) it is difficult to directly validate the *in-vivo* results obtained in this study. However, a number of fundamental relationships between resting physiological parameters have consistently been observed across groups of healthy individuals. Here we compare these observed relationships against the acquired data to infer the relative error and bias for each analysis method. One of the most frequently reported relationships in the healthy human brain is that resting blood flow is linearly correlated with resting oxygen metabolism (Coles et al., 2006; Lebrun-Grandie et al., 1983; Leenders et al., 1990; Powers et al., 2011; Scheinberg and Stead, 1949). Additionally, PET data suggests that the OEF should be approximately uniform across the cerebral grey matter e.g. (Hyder et al., 2016). Thus, we can use the coefficient of variation (COV) of grey matter OEF estimates as an indicator of parameter error, and examine the variation in the slope of the CBF-CMRO_2_ relationship to infer the proportional bias or sensitivity to physiological variation of CMRO_2_ estimates.

As in the simulation experiments we investigated the *in-vivo* analysis for varying levels of regularization in the rNLS analysis and compare this to the ML results. Figure 4b plots the COV in OEF estimates for increasing levels of regularization against the slope of the CBF-CMRO_2_ regression (normalized by the slope of the ML estimate). As predicted by the simulations, the slopes of the ML estimates and the rNLS estimates are similar when little regularization is applied, with the slope of the rNLS estimates slightly reduced compared to the ML approach. As more regularization is applied the COV of OEF estimates is reduced and the slope between CBF and CMRO_2_ decreases, clearly demonstrating the trade-off between variance and bias. Again, as predicted by the simulations, the COV in ML estimates is significantly less than COV in rNLS estimates for a similar CBF-CMRO_2_ slope.

**Figure 4.**
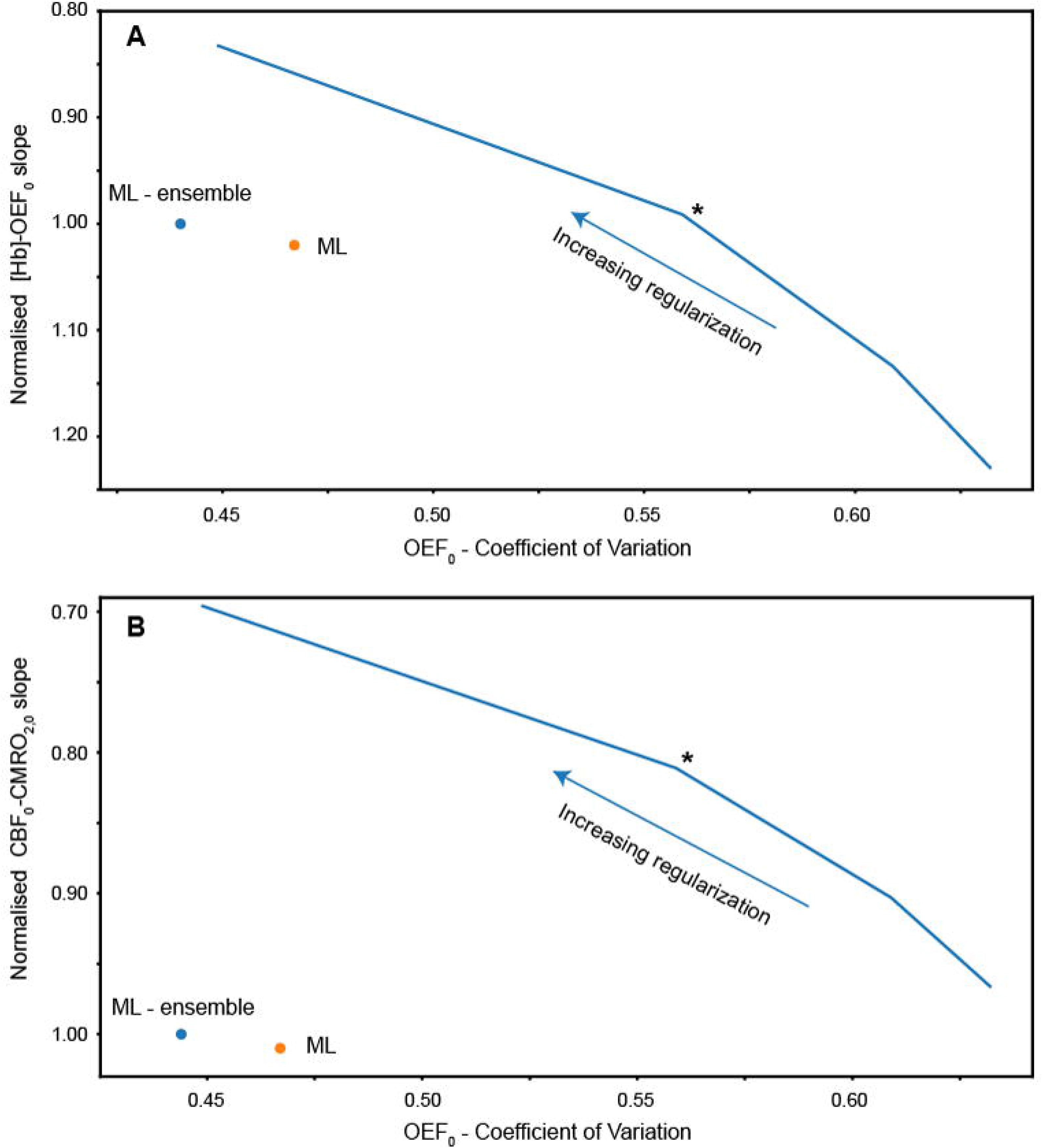
A. Coefficient of variation of grey matter OEF_0_ estimates versus slope of [Hb]-OEF_0_ relationship for each analysis method (rNLS fitting evaluated with increasing levles of regularization). The [Hb]-OEF_0_ slope has been normalised by the ML ensemble estimate of the [Hb]-OEF_0_ slope. B. Coefficient of variation of grey matter OEF_0_ estimates versus the slope of the CBF-CMRO_2_ relationship, normalised by the ML (ensemble) estimate of the CBF-CMRO_2_ slope. Solid blue line plots the coefficient of variation against the slope for increasing regularization weighting for regularized non-linear least squares analysis. The asterisk indicates the chosen level of regularization for subsequent analysis/comparisons.

To investigate the bias in OEF estimates we take advantage of another physiological relationship reported in the literature; cerebral oxygen extraction is inversely related to [Hb] (Ibaraki et al., 2010) and the closely related parameter Hct (Morris et al., 2018). Taking the same approach as before we observe *in-vivo* results that closely match predictions from the simulation, see figure 4a. As in the simulations, the slope in the [Hb]-OEF relationship is similar between the ML method and rNLS approach for a moderate amount of regularization. However, the slope is substantially increased when using minimal regularization, and reduced when applying strong regularization.

Figure 5 shows scatter plots of the grey matter CBF-CMRO_2_ and [Hb]-OEF relationships observed with the ML and rNLS methods across the 30 healthy volunteers studied. The rNLS results are shown for a single level of regularization, where the slope of the [Hb]-OEF relationship most closely matches that of the ML analysis (see figure 4). The coefficient of determination is greater for the ML approach for each relationship, with R^2^ values of 0.56 and 0.35 for the CBF-CMRO_2_ and [Hb]-OEF relationships, compared to 0.34 and 0.14 for the rNLS approach (p<0.05 for all correlations).

**Figure 5.**
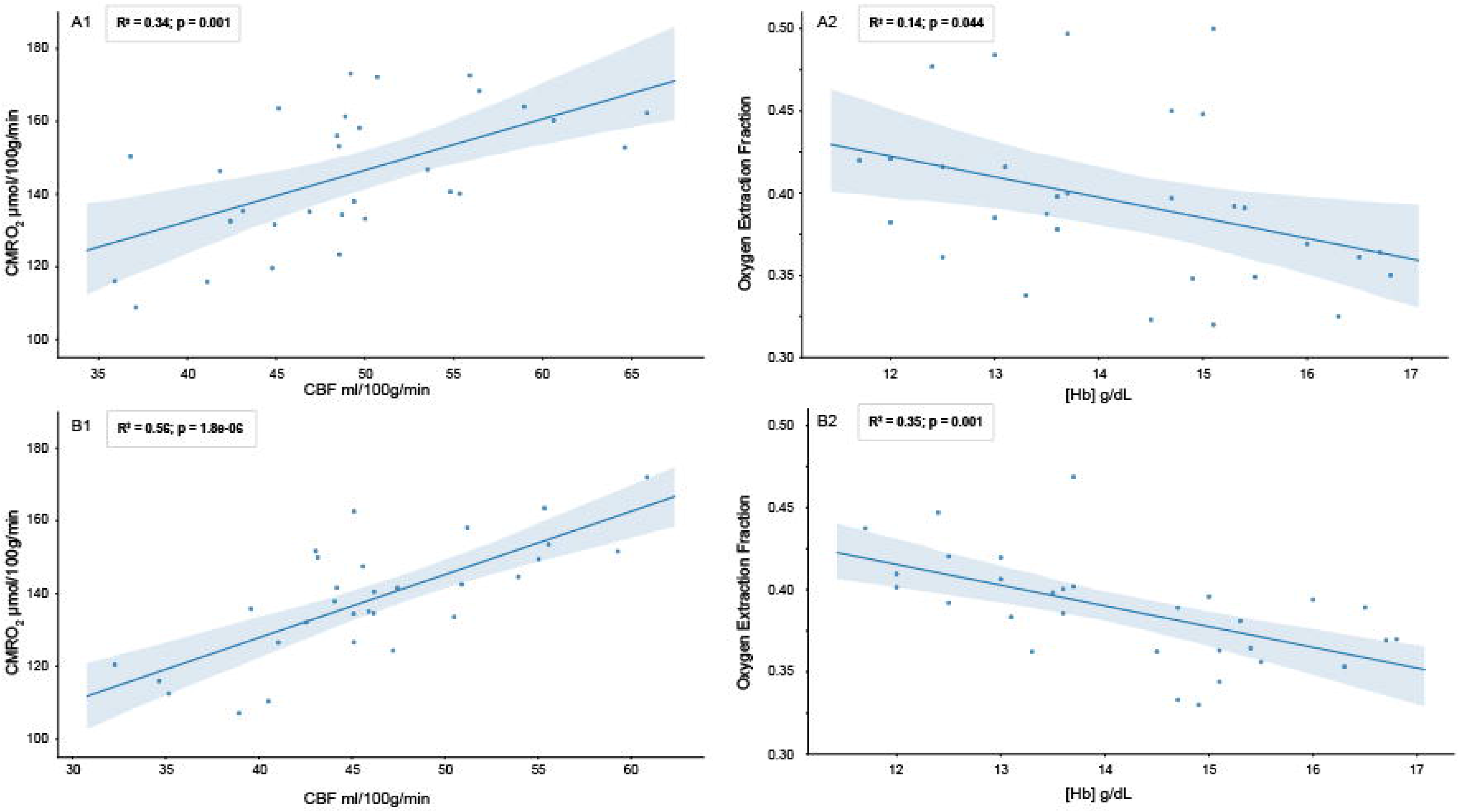
Scatter plots of grey matter CBF-CMRO_2_ and [Hb]-OEF relationships observed with rNLS (A1 and A2)and ML ensemble (B1 and B2) methods across 30 heathy volunteers.

Table 3 reports the results of a bivariate analysis of [Hb] against OEF and CBF for both analysis methods. The slopes of the relationship between OEF and [Hb] are similar to that reported in healthy subjects by (Ibaraki et al., 2010), −1.75 Hb (g/dL). As per Ibaraki et al. the relationship between CBF and OEF did not reach significance (p=0.44) for the ML approach, however a significant negative correlation was observed in the rNLS analysis (p=0.005). A univariate analysis of CMRO_2,0_ against CBF_0_ is consistent with that observed in healthy controls by (Powers et al., 2011) (β1 = 0.2) for both analysis methods, β1 = 0.32 (p<0.001) and β1 = 0.24 (p<0.001) for the ML and rNLS approaches respectively.

**Table 3.** Results of a bivariate regression of [Hb] against CBF_0_ and OEF_0_ grey mater estimates for 30 healthy volunteers analyzed with the ML (ensemble of MLPs) and rNLS fitting methods.

| Predictor | ML $\beta 1$ (p value) | rNLS $\beta 1$ (p value) |
| --- | --- | --- |
| OEF | -1.42 (0.001) | -2.23 (0.001) |
| CBF | -0.07 (0.44) | -0.37 (0.005) |
| Intercept | 61.95 (<0.001) | 89.48 (<0.001) |

Figure 6 shows a comparison between CBF_0_, OEF_0_ and CMRO_2,0_ parameter maps calculated with the ML method (single MLP network and ensemble of 40 networks) and the rNLS method. The image shows 7 slices from a single subject, which have been interpolated for display using cubic b-spline interpolation (Ruijters and Thevenaz, 2012) using FSLeyes (10.5281/zenodo.1470761). As expected OEF_0_ is not well estimated in the white matter, due to the T_1_ decay of the arterial spin labeling signal and the longer arrival time of white matter blood. Across grey matter containing voxels maps of OEF_0_ calculated with the ML methods are more uniform than those calculated with the rNLS approach, with the ensemble approach visibly outperforming the singe network MLP estimates. These observations are consistent with the results of the simulations and the grey matter COV observed for *in-vivo* OEF_0_ estimates. However, it is also apparent from the images that each method demonstrates sensitivity to regional susceptibility effects. For example, in the pre-frontal cortex and inferior temporal lobes the images show greater variability in OEF_0_ estimates, with regions of both over and under-estimation apparent. This instability is likely due to reduced BOLD SNR in these locations and alteration of the susceptibility of air in and around the nasal cavity and paranasal sinuses due to modulation of the inspired oxygen content during data acquisition. It is clear that the ML estimates, in particular those made from the ensemble of MLPs, are more robust to such regional susceptibility effects.

**Figure 6.**
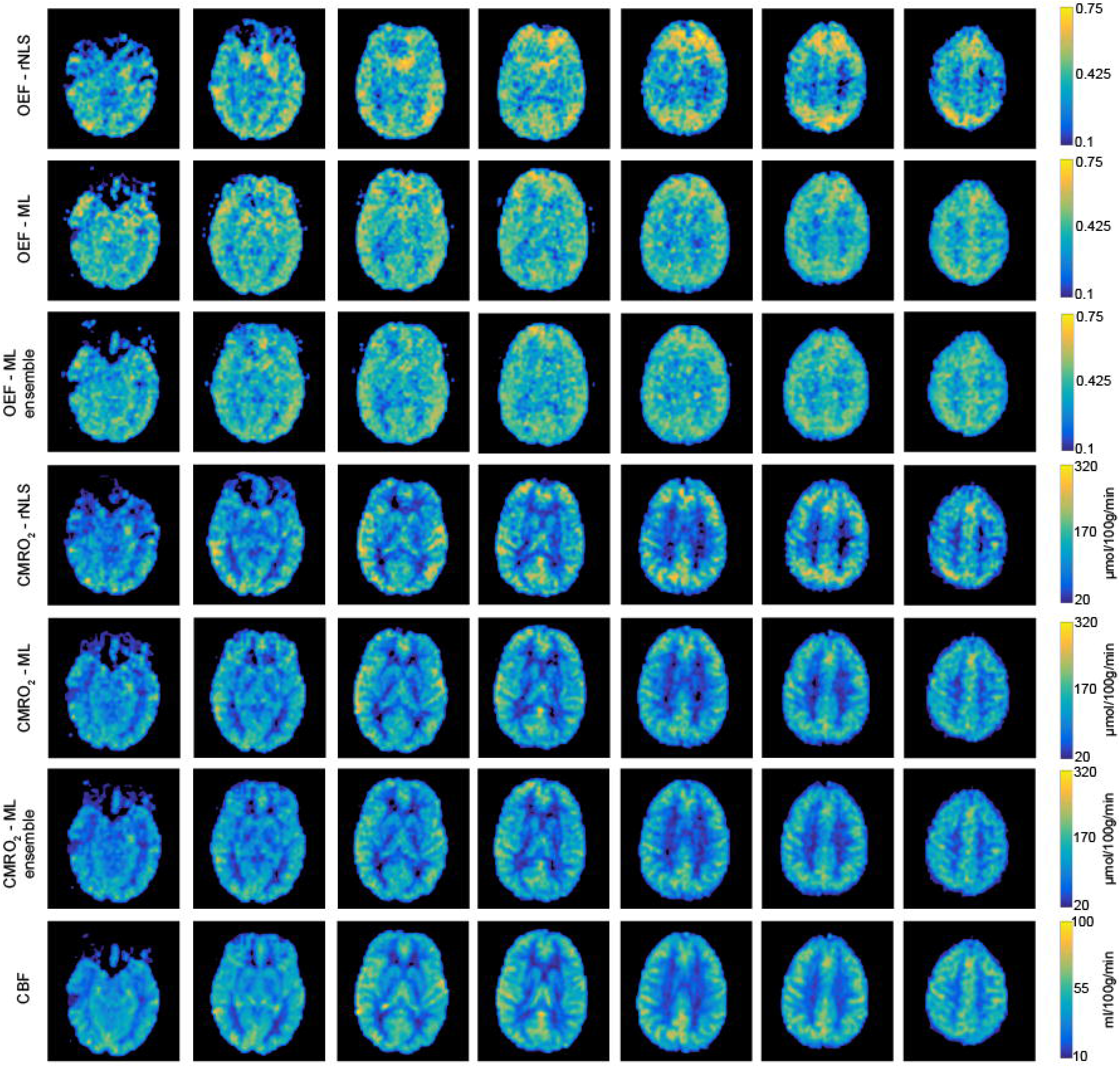
Example parameter maps (CBF_0_, OEF_0_, and CMRO_2,0_) from a single subject for each analysis method. Machine learning estimates of OEF_0_ are more uniform than regularized non-linear least squares estimates. Using an ensemble of MLP networks further reduces the spatial variation in OEF_0_ estimates.

The *in-vivo* analysis also highlights the improvement in computational efficiency of the proposed method. The rNLS approach took approximately 20 minutes to analyze a complete dataset on a standard laptop (2.8 Ghz Intel Core i7, 16GB memory), while the ML approach was able to complete the same analysis in approximately 10 to 20 seconds (depending on the number of networks in the ensemble of MLP regressors).

## 6 Discussion and Conclusions

Instability in parameter estimates made using noisy *in-vivo* data may be reduced by incorporating prior knowledge of physiological parameters, e.g. (Chappell et al., 2010; Frau-Pascual et al., 2014; Germuska et al., 2016; Mesejo et al., 2015). Previous investigation of such methods (Germuska et al., 2016) suggests that they are an effective means to increase the robustness of CMRO_2_ estimates made with dc-fMRI. However, these methods are computationally expensive and must necessarily make a trade off between parameter uncertainty and parameter sensitivity. Thus, they are not well suited to high throughput or rapid data analysis and care must be taken when using such methods not to unduly bias parameter estimates towards the priors. In the work presented here we take a different approach by training a machine learning implementation that is robust to input noise. Given an appropriately selected (or generated) training dataset, a well-implemented solution will be unbiased, robust, and have a low computational overhead.

Computer modeling suggests that the proposed method outperforms previous analysis methods both in terms of uncertainty and bias. *In-vivo* data supports the predicted improvement in uncertainty with a significant reduction in the COV of grey matter OEF_0_ estimates when compared to a regularized non-linear least squares fitting of the data. Additionally, agreement was found between the predicted behaviors of each method and their associated biases when compared to reported physiological relationships. Qualitatively, the *in-vivo* parameter maps suggest that the ML approach, especially when paired with an ensemble implementation, is more robust to physiological noise; producing physiologically plausible parameter estimates in challenging brain regions, e.g. near the frontal sinuses. Such physiological noise was not modeled in the training data so it is perhaps unexpected that the ML method is robust to these noise sources. However, it is plausible that the discriminative features identified from the frequency-domain representation of the data during training are less sensitive to these regional susceptibility changes than a traditional time-domain fit of the data. It is possible that this aspect of the ML approach could be enhanced by extending the training data to include such regional susceptibility changes, either on their own or in combination with a spatially informed approach to data fitting.

The use of an ensemble of MLP networks reduced parameter uncertainty in simulation and reduced the coefficient of variation in grey matter OEF_0_ estimates *in-vivo*, demonstrating its utility in this application. However, it is anticipated that enforcing network diversity during training could make further improvements in performance. As it is has previously been demonstrated that, in the presence of noise, the performance of an ensemble of networks can always be improved by explicitly encouraging diversity during training (Reeve and Brown, 2018).

The machine learning implementation presented here employs a combination of proven signal processing (time-frequency transformation) and machine learning methods (decision trees and fully connected artificial neural networks) that have been shown to select appropriate features for learning and are robust to input noise. The proposed analysis pipeline demonstrates an improvement in both the accuracy and precision in parameter estimates compared to published methods, and is appropriate for the study of both healthy volunteers and in clinical investigations. However, there are still many avenues that could be explored both in terms of signal processing and machine-learning. For example time domain data could be converted to 2D time-frequency representations such as a spectrogram, or into spectrogram-like representations using wavelet transforms (for increased time resolution). This type of pre-processing would open the door to the application of 2D convolution neural networks (CNN) that have been so successfully applied in the domain of image processing. It is possible that the application of such approaches could further improve the performance of machine learning when analyzing dc-fMRI data. However, a thorough investigation of all available machine learning methods and associated pre-conditioning of the data is beyond the scope of the current study, which focuses instead on the realization of a practical solution by combining well-proven techniques for the analysis of signal data.

All *in-vivo* analysis in this manuscript is performed in the absence of spatial smoothing, which is often employed to improve statistical estimates made from fMRI data (Friston et al., 1995). We chose not to employ spatial smoothing in this analysis for two principle reasons: first any such spatial filtering implies a prior assumption regarding the spatial extent of any variation (Rosenfeld and Kak, 1982), and can thus lead to unwanted loss of sensitivity to physiological variation; second we did not want to increase the potential contamination of grey matter voxels with non-tissue signals, such as CSF or macrovessels (both of which are not included in the underlying signal model). The current study does not make any direct comparison between smoothed and unsmoothed analysis pipelines, however the presented method clearly avoids any possible smoothing artefacts that might otherwise bias the analysis.

A limitation of the proposed method is the need to train new regressors for a given gas paradigm and set of acquisition parameters, e.g. arterial spin labeling tagging duration, repetition time and duration of the acquisition. In addition, there is a requirement that the *in-vivo* gas manipulation does not deviate significantly from the range of simulated designs. While it is a relatively straightforward process to retrain the regressors with a new set of parameters, to match the local acquisition protocol, the scope of the method could be increased if individualized gas traces could be incorporated into the training data; allowing a single pre-trained implementation to be applied across studies.

The simulations and *in-vivo* results suggest that the proposed analysis method could significantly increase the utility of dc-fMRI, reducing the number of participants needed to detect a group difference in oxygen metabolism or oxygen extraction fraction and offering more physiological interpretability of metabolic differences or alteration due to a stimulus. In addition, the significant reduction in processing time and the improved robustness of the individual parameter maps reduces two of the hurdles restricting clinical implementation of such techniques.

## Supporting information

Demographics

## 8 Author Contributions

MG. Wrote the manuscript, developed and implemented the methods, and analysed the in-vivo data. HC. Acquired and processed data and edited the manuscript.

TO. Created and provided code used in the prototype pseudo-continuous arterial spin labeling pulse sequence.

FF. Assisted in the implementation of the prototype arterial spin labeling pulse sequence.

VT. Chief investigator overseeing study design and data collection for a subset of healthy controls.

KM. Principal investigator overseeing study design and data collection for a subset of healthy controls.

RW. Principal investigator overseeing study design and data collection for a subset of healthy controls

All authors reviewed and edited the manuscript prior to submission.

## 10 Acknowledgments

We would like to thank Wellcome for supporting this work: Wellcome Strategic Award, ‘Multi-scale and multi-modal assessment of coupling in the healthy and diseased brain’, grant reference 104943/Z/14/Z.

## Data Availability Statement

The python code for the machine learning implementation proposed in this manuscript can be found in the fml_pMRI repository https://zenodo.org/badge/latestdoi/189416118. We do not have ethical consent to make the *in-vivo* datasets acquired for this study publically available.

