## Supplementary figures and images for "A frequency-domain machine learning method for dual-calibrated fMRI mapping of oxygen extraction fraction (OEF) and cerebral metabolic rate of oxygen consumption (CMRO_2_)"

### Demographics

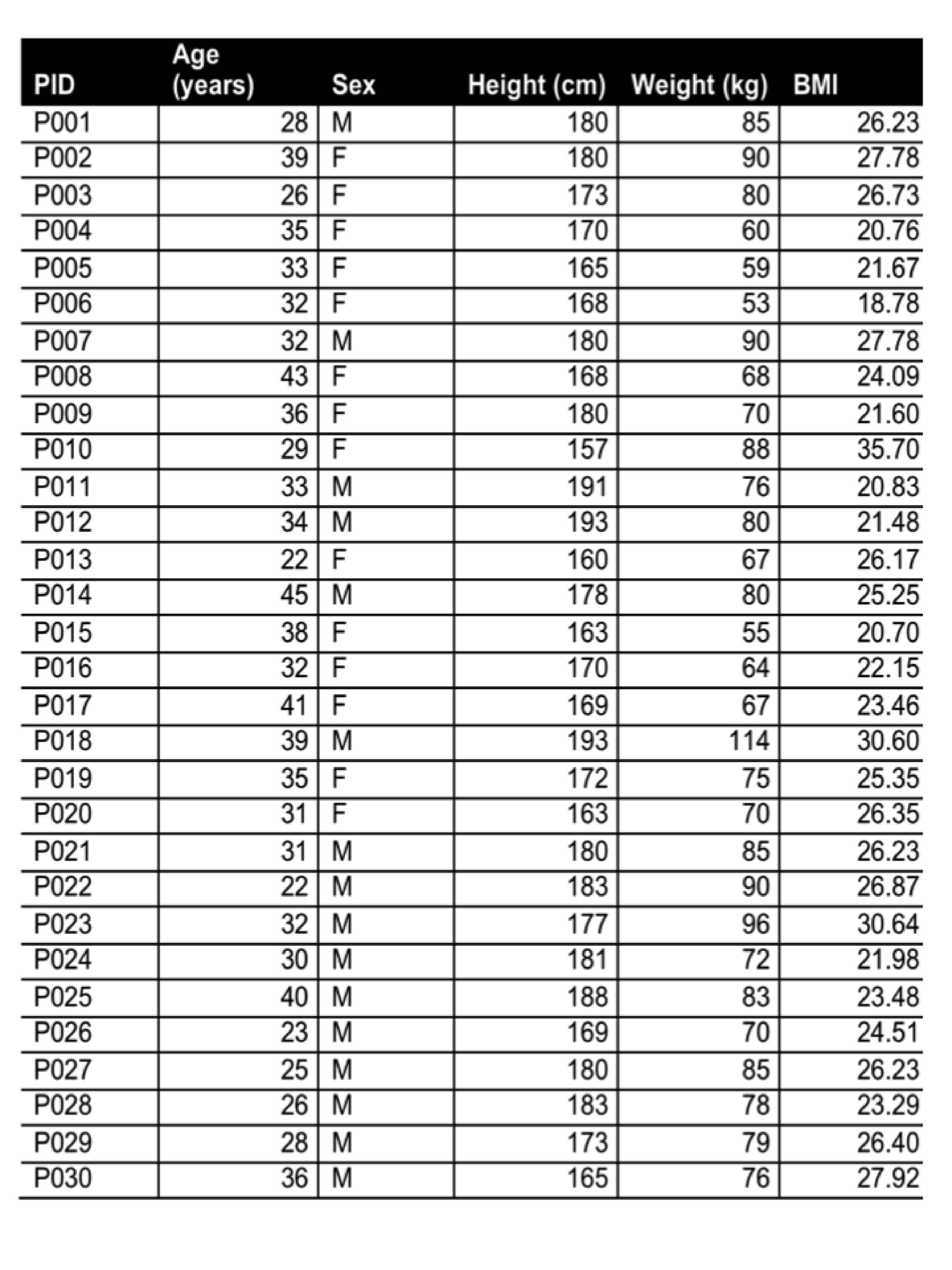
